# The Drosophila *attP40* docking site and derivatives are insertion mutations of *MSP300*

**DOI:** 10.1101/2022.05.14.491875

**Authors:** Kevin van der Graaf, Saurabh Srivastav, Pratibha Singh, James A McNew, Michael Stern

## Abstract

The ϕC31 integrase system is widely used in *Drosophila* to allow transgene targeting to specific loci. Over the years, flies bearing any of more than 100 *attP* docking sites have been constructed. One popular docking site, termed *attP40*, is located close to the *Nesprin-1* orthologue *MSP300* and lies upstream of certain *MSP300* isoforms and within the first intron of others. Here we show that *attP40* causes larval muscle nuclear clustering, which is a phenotype also conferred by *MSP300* mutations. We also show that flies bearing insertions within *attP40* can exhibit decreased *MSP300* transcript levels in third instar larvae. Finally, chromosomes carrying certain “transgenic RNAi project” (TRiP) insertions into *attP40* can confer pupal or adult inviability, or infertility. These phenotypes do not require transcription from the insertions within *attP40*. These results demonstrate that *attP40* and insertion derivatives act as *MSP300* insertional mutations. These findings should be considered when interpreting data from *attP4*0-bearing flies.

## Introduction

For several years, Drosophila investigators have used a genome integration method based on the site-specific ϕC31 integrase (Thorpe and Smith 1998) to target transgenes to specific loci (Groth 2004). With this method, ϕC31 integrase catalyzes sequence-directed recombination between a phage attachment site (*attP*, present within each of >100 *attP* “docking sites” in Drosophila) and a bacterial attachment site *(att*B, present within the integrating plasmid) (Thorpe *et al*. 2000; Groth and Calos 2004; Bateman *et al*. 2006; Venken *et al*. 2006; Bischof *et al*. 2007). By allowing transgene insertion into specific, defined, docking site sequences, the ϕC31 integrase method increases the reproducibility and decreases the variability of transgene expression observed with the random transgene integration utilized by P elements.

Two docking sites, *attP2*, located at position 68A4 on chromosome III and *attP40*, located at position 25C on chromosome II, are widely used docking sites for LexA drivers and Gal4-driven Transgenic RNAi Project (TRiP) insertions (Zirin *et al*. 2020). These two *attP* docking sites are favorable because they express inserted transgenes at high levels while maintaining low basal expression (Perkins *et al*. 2015). In fact, the Drosophila stock center at Bloomington, IN, reports possessing 16,503 Drosophila lines carrying *attP40* and 14,970 lines carrying *attP2*; most lines carry TRiP insertions or the activation domains or DNA binding domains from the Janelia split-Gal4 collection (Annette Parks, personal communication). Although originally reported to be located in an intergenic region, between *CG14035* and *MSP300* (Markstein *et al*. 2008), *attP40* lies within the first intron of certain *MSP300* isoforms ((Larkin *et al*. 2021) FlyBase FB2022_02). This observation raises the possibility that *attP40* might act as an insertional mutation for *MSP300*. Indeed, it was previously reported that certain insertions into *attP40* could cause spreading of the H3K27me3 mark over the large *MSP300* exon (De *et al*. 2019). Thus, transgenes inserted within the *attP40* docking site might affect expression of at least a subset of *MSP300* isoforms.

MSP300 (Muscle-specific protein 300 kDa) is a nuclear-associated Nesprin1 orthologue and a component of the Linker of Nucleoskeleton and Cytoskeleton (LINC) complex (Kim et al., 2015; Volk, 1992). The C-terminal domain contains a Klarsicht/Anc1/Syne Homology (KASH) domain that interacts with Sad1p/UNC-84 (SUN)-domain-containing proteins, connecting the outer and inner nuclear membranes (Xie and Fischer, 2008; McGee et al., 2006). In Drosophila larvae, *MSP300* transcription has been reported in muscle (Volk, 1992) and fat body (Zheng *et al*. 2020). In larval muscle, MSP300 forms striated F-actin-based filaments that lie between muscle nuclei and postsynaptic sites at the neuromuscular junction. MSP300 also wraps around immature boutons in response to electrical activity and is required for postsynaptic RNA localization and synaptic maturation (Packard *et al*. 2015). MSP300 is also required for normal nuclear localization in muscle cells and for integrity of muscle cell insertion sites into the cuticle (Volk, 1992; Volk, 2013; Zhang et al., 2010). MSP300 isoforms lacking the KASH domain confer deficits in larval locomotion, localization of the excitatory neurotransmitter receptor GluRIIA at the neuromuscular junction (NMJ), and proper NMJ functioning, independently of its role in muscle nuclear positioning (Morel *et al*. 2014). Non-muscle deficits conferred by *MSP300* mutations include defects in oocyte development and female fertility (Yu *et al*. 2006). In humans, mutations in the Nesprin family are associated with several musculoskeletal disorders, including bipolar disorder, autosomal recessive cerebellar ataxia type 1 (ARCA1), X-linked Emery-Dreifuss muscular dystrophy (EDMD) and are risk factors for schizophrenia and autism (Rajgor and Shanahan 2013).

Here, we show that flies carrying *attP40* exhibits a nuclear clustering phenotype in larval muscle, which suggests that *attP40* is an *MSP300* insertional mutation. Further, we show that inserting transgenes into *attP40* can increase severity of this phenotype. We use quantitative RT-PCR (Q-PCR) to show that insertions within *attP40* decrease *MSP300* transcript levels in 3^rd^ instar larvae. Finally, we show that chromosomes carrying certain TRiP insertions constructed from the *Valium 20* vector (Perkins *et al*. 2015) into *attP40* confer recessive lethality or sterility. Because of the variable effects of different transgene insertions into *attP40*, investigators should use caution in interpreting data collected from *Drosophila* carrying these insertions.

## Materials and Methods

### Drosophila stocks

All fly stocks were maintained on standard cornmeal/agar *Drosophila* media: 69.1 g/l corn syrup, 9.6 g/l soy flour, 16.7 g/l yeast, 5.6 g/l agar, 70.4 g/l cornmeal, 4.6 ml/l propionic acid and 3.3 gm/l nipagin. Flies carrying *attP2* and *attP40* lacking insertions were retrieved as white-eyed progeny from transgene insertions carried out at GenetiVision (Houston, TX). The Drosophila Stock Center at Bloomington, Indiana provided TRiP *JNK* (#57035), TRiP *spatacsin (spat)* (#64868), TRiP *Rop (#51925)*, TRiP *Spartin (#37499)*, TRiP *atlastin (atl)* (#36736), TriP *Mcu* (#42580), *13XLexAop2-IVS-myr::GFP* (#32210), *LexA::Mef2* (#61543), *CyO-TbA* (*#36335*) and *LexA::nSyb* (#52817). All experiments were performed on *Drosophila* that had been reared and maintained at room temperature (22°C) with a 12h: 12h light:dark cycle unless otherwise indicated.

### Immunocytochemistry

All larvae were dissected in HL3.1 (Feng *et al*. 2004) in a magnetic chamber, and fixed in 4% paraformaldehyde for 10 minutes, then washed in PBS with 0.3% Triton-X (PBS-T) and blocked for 30 minutes in PBS-T with 1% BSA. Samples were incubated overnight at 4°C with primary antibody, washed thoroughly with PBS-T and then incubated for 2.5 hours at room temperature with secondary antibody. Samples were then washed with PBS-T and mounted in VectaShield Antifade Mounting Medium containing DAPI (Vector laboratories; H-1200-10).

Alexa Fluor® 647 phalloidin (1:200) was used to visualize F-actin. All images were acquired on a Zeiss LSM800 with an Airyscan confocal microscope.

### Nuclear clustering analysis

Third instar larval body wall muscle 6 was chosen for all nuclear clustering analysis. Images were opened in ImageJ and nuclei clusters in muscle 6 were counted. We defined a “cluster” as two or more nuclei in which the distance between two nuclear borders was less than five microns. We analyzed six larvae from hemisegments A2-A4 (18 hemisegments total) for each genotype.

Microsoft excel was used to import all nuclear clustering data. “Normal” muscles contained no clusters. For muscles that contained cluster(s), we determined cluster number and nuclei number per cluster. Cluster sizes for each genotype were plotted on a frequency histogram. The percentage of normal hemisegments for each genotype was plotted on a column graph. For each hemisegment we calculated the percentage of nuclei in clusters using the following calculation: number of nuclei within clusters divided by total number of nuclei in muscle 6 multiplied by 100. Then, the average of three hemisegments per larva was calculated and plotted on a column graph. Data was visualized and analyzed using GraphPad Prism v9.3.1 or Synergy Software KaleidaGraph v4.5.2.

### Quantitative RT-PCR (Q-PCR)

Primers were designed with PrimerBLAST software according to the published sequence of *MSP300*. We amplified and analyzed two *MSP300* regions: first, the 3’ end region, which amplifies transcripts *RD, RG, RH, RI, RJ, RK, RL* and *RM* (transcripts that contain the KASH domain), and second, the middle region, which amplifies transcripts *RB* and *RF* (*B/F*) (transcripts that lack the KASH domain) (Figure 1). For the 3’ end region, we used the forward primer sequence 5’-TCAACCTCTTCCAATGCAGGC-3’, and the reverse primer sequence 5’-CGCCAGAACCGTGGTATTGA-3’. The *B/F* forward primer sequence was 5’-CACGTACTTGCCGCACGAT-3’, and *B/F* reverse primer sequence was 5’-ATTTTTGACACGTTCCCGGC-3’. For amplification of *Rp49*, chosen as the housekeeping gene, the forward primer sequence was 5’-TGTCCTTCCAGCTTCAAGATGACCATC-3’ and the reverse primer sequence was 5’-CTTGGGCTTGCGCCATTTGTG-3’. Total RNA (500 ng) was extracted from frozen whole larvae and larval fillets with Direct-zol™ RNA MicroPrep (Zymo Research) according to manufacturer’s protocol. The yield of RNA was estimated with the Nanodrop2000 (ThermoFisher). The A_260_:A_280_ ratio was between 1.8-2.1. Superscript™ III First-Strand Synthesis System (ThermoFisher) was used to generate cDNA according to manufacturer’s protocol. Reverse transcribed cDNA was then amplified in a 20 µl PCR reaction by the ABI Prism 7000 system (Applied Biosystems) with the universal conditions: 50°C for 2 min, 95°C for 10 min, and 40 cycles (15s at 95°C, 1 min at 60°C). Each sample contained 10 whole larvae. Three separate biological samples were collected from each genotype, and triplicate measures of each sample were conducted for amplification consistency. Data were analyzed with the relative 2^- ΔΔCt^ method to ensure consistency (Livak and Schmittgen 2001).

**Figure 1.**
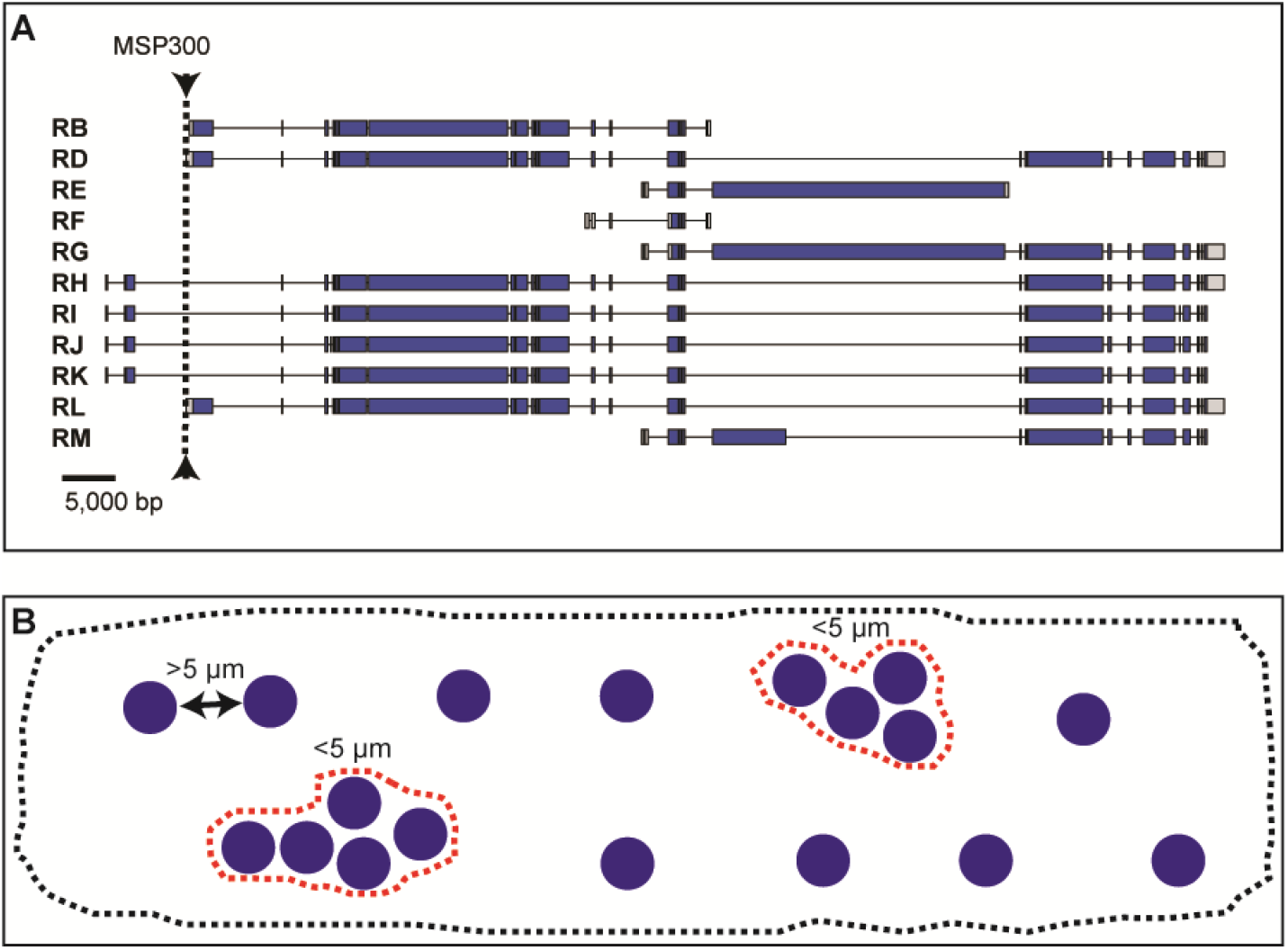
Overview of MSP300 transcripts and nuclear clustering measurements. A) Overview of the 11 annotated MSP300 transcripts in *Drosophila melanogaster*. Arrowhead/dotted line indicates location of the *attP40* docking site. Predicted exons shown with blue bars (translated regions) or gray bars (5’ UTR or 3’ UTR), introns with thin lines. B) Schematic representation of nuclear clustering in larval body wall muscle 6. If borders between two nuclei are less than 5 µm apart it is counted as a cluster (dotted red lines). Dotted gray line outlines muscle 6.

### Pupal size measurements

We measured length and width in pupae homozygous for each of six TRiP lines and the *attP40* parent chromosome. Pupal length was determined from top to bottom, excluding the anterior and posterior spiracles. Pupal width was determined by measurement at the midpoint along the pupal anterior/posterior axis.

### Viability measurements

Each of six TRiP lines were placed in combination with balancer “*CyO-TbA”* (Lattao *et al*. 2011), which carries the *Tb*^*1*^ dominant “Tubby” marker and enabled us to unambiguously genotype balancer-containing from balancer-lacking larvae, pupae and adults. Each of these six TRiP lines were then brother-sister mated and F1 progeny reared in uncrowded vials. The number of Tubby and non-Tubby pupae were counted. Non-Tubby pupae were removed and placed into fresh vials. After seven days, the number of successful eclosions was counted. To monitor viability in the control *attP40* line, pupae reared in uncrowded vials were collected and after seven days the number of successful eclosions was counted.

### Fertility measurements

We measured male and female fertility in adults homozygous for each of six TRiP lines and the *attP40* parent chromosome. To measure male fertility, single males were placed in vials with two phenotypically wildtype and fertile virgin females, and ability to generate larval progeny was measured. To measure female fertility, single females were placed in vials with two phenotypically wildtype and fertile males, and ability to generate larval progeny was measured.

### Construction of *Aop-atlRNAi*

We chemically synthesized the reported sequence of the TRiP *atl* short hairpin with Xho1 and *Xba1* sites added to the left and right ends, respectively. (TCGAGAGTCTGGTATAGGTCATTAGTTTAtagttatattcaagcataTAAACTAATGACCTATACC AGGCT – lower case letters indicate the loop sequence). This construct was introduced into the *Xho1-Xba1* sites of *pJFRC19-13XlexAop2* (Addgene, Inc) and introduced into embryos at the *attP40* site using with ϕC31-mediated recombination (GenetiVision, Houston TX). Insert-containing flies were recognized by eye color.

### Statistical analysis

For percentage of nuclei in clusters analysis, a One-way ANOVA with Tukey post hoc test was performed. For pupa length and width analysis, a One-way ANOVA with Bunnett post hoc test was performed. All statistical tests were performed in GraphPad Prism v9.3.1.

### Availability

Fly stocks are available upon request. The authors affirm that all data necessary for confirming the conclusions of the article are present within the article and figures.

## Results

### The *attP40* docking site is located within or upstream of *MSP300*, depending on the isoform

*attP2* and *attP40* are two widely used *attP* docking sites in *Drosophila* for insertions of LexA drivers, Gal4-driven TRiP insertions, and other constructs. Both docking sites provide high levels of induced transgene expression while maintaining low basal expression. As of February 2022, the Bloomington Drosophila Stock Center (Bloomington, IN) provides 16,503 stocks carrying *attP40* and 14,941 stocks carrying *attP2* (Annette Parks, personal communication). The *attP40* chromosomal location is closest to *MSP300*, which expresses 11 different isoforms. *attP40* lies within intron 1 for transcripts RH, RI, RJ and RK and upstream of transcripts RB, RD, RE, RF, RG, RL and RM (Figure 1A). These results raise the possibility that *attP40* could affect *MSP300* transcript levels. Indeed, De et al. (De *et al*. 2019) reported that certain transgene insertions into *attP40* alter the H3K27me3 epigenetic mark over at least a part of *MSP300*. Similarly, in a Ph.D. thesis, (Cypranowska 2020) showed that *attP40* decreased *MSP300* transcription in certain genetic backgrounds and conferred synaptic phenotypes at the larval neuromuscular junction consistent with decreased MSP300 activity.

Given previous reports that *MSP300* variants alter muscle myonuclear spacing (Elhanany-Tamir et al., 2012; Volk, 1992), we hypothesized that effects of *attP40* on *MSP300* expression might alter larval muscle nuclear clustering. To address this hypothesis, we first defined a “cluster” as two or more nuclei in which the distance between two nuclear borders was less than 5 µm (Figure 1B). Then, for every nucleus in a larval body wall muscle, we measured the distance to its nearest neighbor, using larval body wall muscle 6 as assay platform. Using this approach, we observed abnormalities in nuclear positioning in larvae homozygous for *attP40* vs. control larvae (homozygous for *attP2*) or larvae heterozygous for *attP40* and *attP2* (Figure 2A-C). In particular, larvae homozygous for *attP40* exhibited more nuclear clusters and clusters of greater size than control or heterozygous larvae (Figure 2D), and a significantly greater number of nuclei within clusters (Figure 2E). As a result, larvae homozygous for *attP40* exhibited fewer hemisegments with a normal nuclear distribution than control or heterozygous larvae (Figure 2F). We conclude that *attP40* causes a recessive muscle nuclear clustering phenotype.

**Figure 2.**
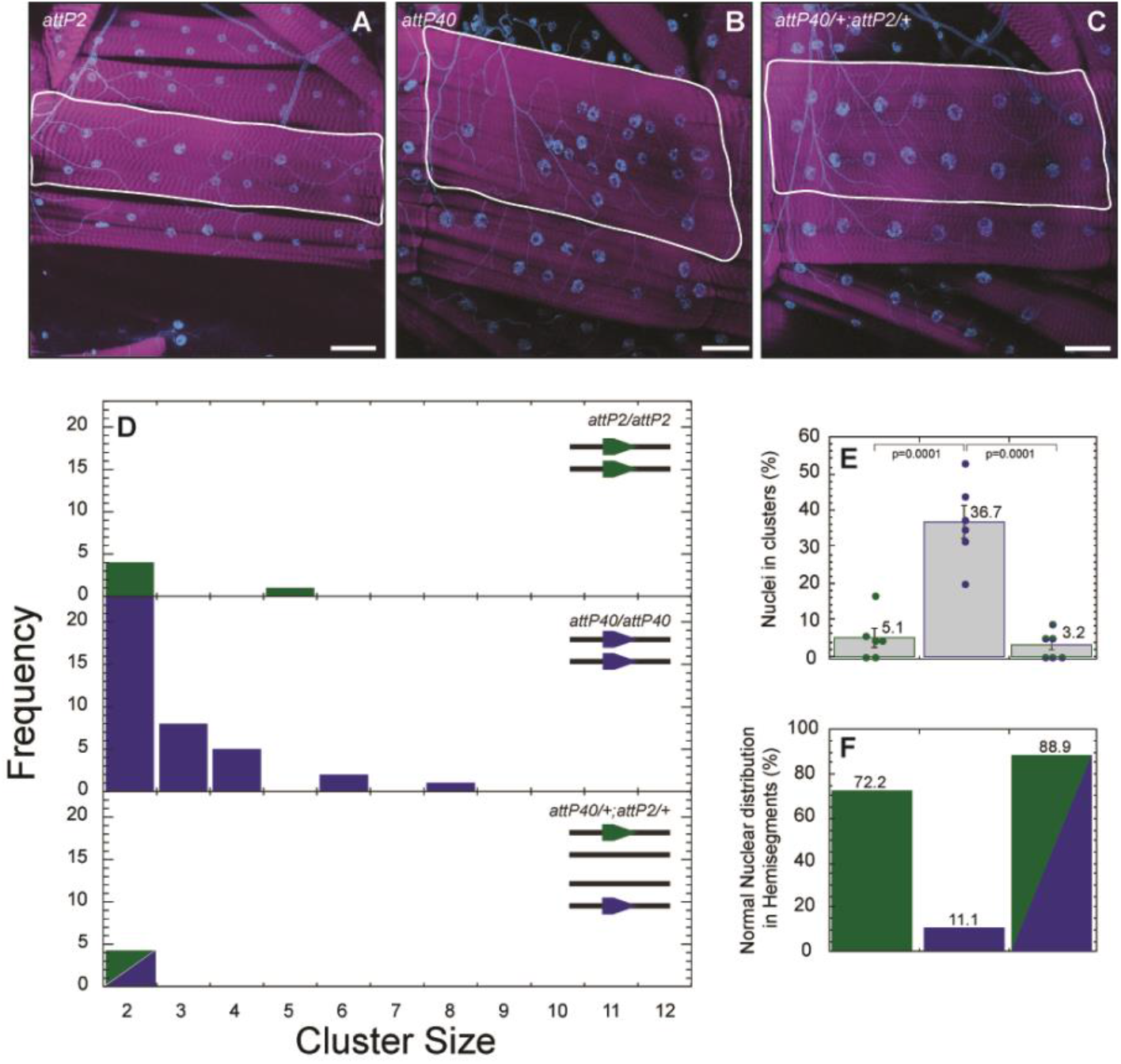
Muscle nuclei clustering phenotype observed in *attP40* larvae. A-C) Representative images of nuclei positioning within muscle 6 for larvae carrying *attP2* (A), *attP40* (B) and *attP40*/+; *attP2*/+ (C). Nuclei are labeled with DAPI (blue), and muscle actin is labeled with phalloidin (magenta). Muscles are outlined with white lines. D) Frequency distribution of the number of nuclei within each cluster within muscle 6 for each genotype shown in upper right. Lines in upper right depict the genotype of each homologue; green pentagons indicate the *attP2* docking site, blue pentagon depicts *attP40* docking site E) Percentage of nuclei found within clusters for each genotype. Means +/-SEMs are shown. The following calculation was used: ((number of nuclei within clusters / total number of nuclei) x 100). F) Percentage of hemisegments with normal nuclear distribution within muscle 6. *attP2* is represented in green, *attP40* in blue and *attP40*/+; *attP2*/+ in half green/half blue for panels D-F. For each genotype 3 hemisegments per larvae for 6 larvae total were analyzed. One way ANOVA with Tukey post-hoc test was used for statistical analyses. Scale bar = 50 µm.

### Effects of inserting transgenes into *attP40* on phenotypic severity

To determine if insertions into *attP40* could affect nuclear clustering, we used the functionally neutral *LexAop2-IVS-myr::GFP* reporter introduced into *attP40* and crossed these flies with *attP2, attP40, lexA::Mef2* or *lexA::nSyb*. Representative images of nuclei clustering in body wall muscle 6 for each genotype is shown in Figure 3A-D. First, we found that *LexAop2-IVS-myr::GFP/+* larvae (only one copy of *attP40*) showed only 6.18% of nuclei in clusters and a mostly normal nuclear distribution (Figure 3E-G), indicating that *LexAop2-IVS-myr::GFP*, like “empty” *attP40* (*attP40* lacking an insertion), is recessive. However, when *LexAop2-IVS-myr::GFP* was in combination with empty *attP40*, we found extremely large nuclear clusters (containing up to 12 nuclei; Figure 3E), which were not observed in larvae homozygous for *attP40* (Figure 2D). In addition, 46.4% of nuclei were in clusters compared to 36.7% in larvae homozygous for *attP40* homozygous (Figures 3F and 2E). Thus, transgene insertion into one *attP40* site increases nuclear clustering severity. To determine if inserting transgenes into both *attP40* sites would affect nuclear clustering, we examined nuclear clustering in *LexAop2-IVS-myr::GFP/LexA::nSyb* larvae and found nuclear clustering similar to larvae homozygous for empty *attP40* (Figures 3E and 2D). These results suggest that inserting transgenes into a second *attP40* site has little additional effect on nuclear clustering.

**Figure 3.**
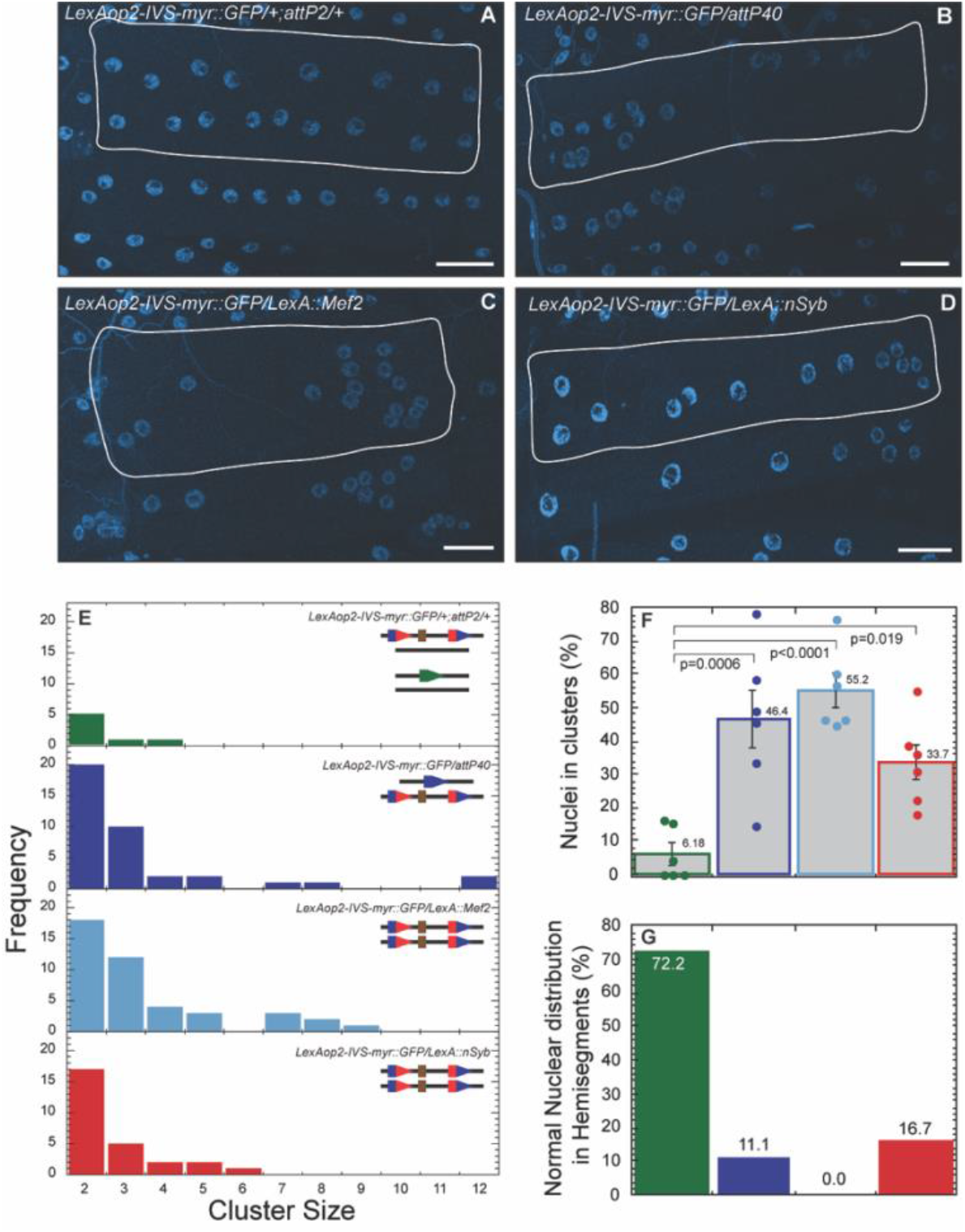
Effects of inserting AOP/LexA transgenes into *attP40* on nuclear clustering severity. A-D) Representative images of nuclei position within muscle 6 for LexAop-IVS-myr::GFP in combination with *attP2* (A), *attP40* (B), *LexA::Mef2* (C) and *LexA::nSyb* (D). Nuclei are labeled with DAPI (blue). Muscles are outlined with solid white lines. E) Frequency distribution of the number of nuclei within each cluster within muscle 6 for each genotype indicated in upper right. Lines in upper right show genotype of each chromosome II homologue (in top right histogram, both chromosome II and chromosome III are shown). Green pentagon represents *attP2*, blue pentagons represent *attP40*, red/blue composite pentagons represent *attP40* carrying indicated insertion. Brown rectangle represents insertion. F) Percentage of nuclei found within clusters for each genotype. Means +/-SEMs are shown. The following calculation was used: ((number of nuclei within clusters / total number of nuclei) x 100). G) Percentage of hemisegments with normal nuclear distribution within muscle 6. In panels E-G, each genotype is represented with a different color as follows: *LexAop2-IVS-myr::GFP/attP2* (green), *LexAop2-IVS-myr::GFP/attP40* (blue), *LexAop2-IVS-myr::GFP/LexA::Mef2* (light blue) and *LexAop2-IVS-myr::GFP/LexA::nSyb* (red). For each genotype 3 hemisegments per larvae for 6 larvae total were analyzed. One way ANOVA with Tukey post-hoc test was used for statistical analyses. Scale bar = 50 µm.

The transgenes studied above did not include a transcriptional “driver” and were unexpressed in the body wall muscle. Given the previous report (De *et al*. 2019) that certain transgene insertions into *attP40* could generate epigenetic marks that spread into *MSP300*, potentially altering *MSP300* expression, we determined if expressing transgenes in muscle would affect nuclear clustering. Thus, we created *LexAop2-IVS-myr::GFP/LexA::Mef2* larvae, in which transgenes in each *attP40* site would be expressed in muscle. We found that these larvae exhibited a more severe nuclear clustering phenotype even than *LexAop2-IVS-myr::GFP/*empty *attP40* (Figure 3E-G), and was the only genotype tested in which every muscle exhibited some nuclear clustering (Figure 3G). These observations suggest that transcribing insertions in *attP40* might further increase the severity of nuclear clustering phenotype.

To determine effects of Gal4-regulated, rather than LexA-regulated, transgenes inserted into *attP40*, we crossed *attP2* or empty *attP40* flies to flies carrying the Gal4-driven shRNA construct targeting *spatacsin (spat)* (CG13531) created by the transgenic RNAi project (TRiP) (Perkins *et al*. 2015). Representative images are shown in Figure 4A-C. We found that both TRiP *spat*/empty *attP40* or homozygous TRiP *spat* exhibited a strong nuclear clustering phenotype (Figure 4D-F) – in fact, larvae homozygous for TRiP *spat* exhibited clusters up to 15 nuclei in size (Figure 4D). However, TRiP *spat/+;attP2/+* larvae exhibited wildtype nuclear positioning, indicating that TRiP *spat* is recessive. Thus, this Gal4-driven transgene behaves in a manner similar to the LexA transgenes described above.

**Figure 4.**
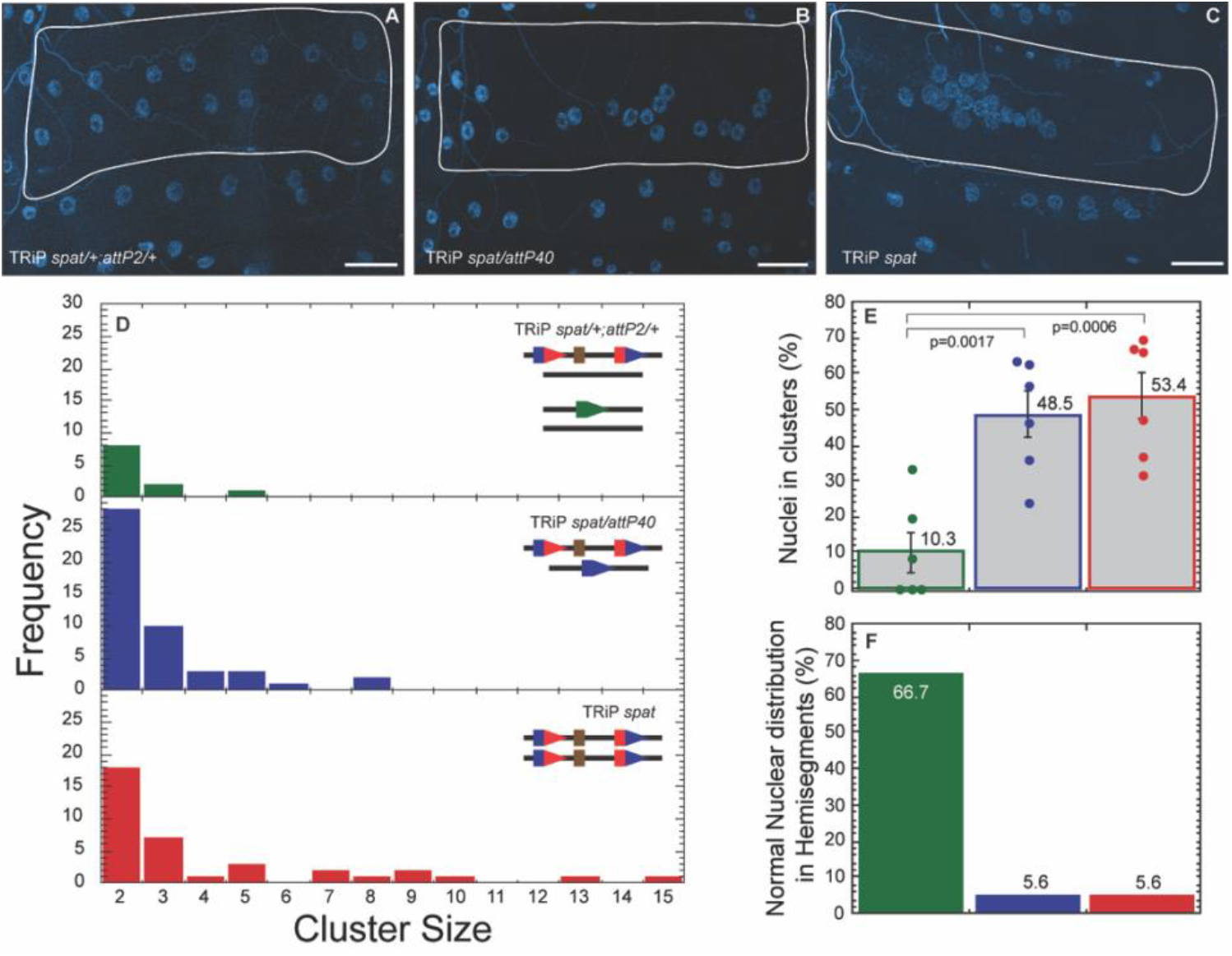
Effects on nuclei clustering of inserting Gal4-regulated TRiP *spatacsin (spat)* into *attP40*. A-C) Representative images of nuclear position within muscle 6 for TRiP *spat*/+; *attP2*/+ (A), TRiP *spat*/*attP40* (B) and TRiP *spat* (C). Nuclei are labeled with DAPI (blue). Muscles are outlined with solid white lines. D) Frequency distribution of the number of nuclei within each cluster within muscle 6 for each genotype indicated in upper right. Lines in upper right show genotype of each chromosome II homologue (in top right histogram, both chromosome II and chromosome III are shown). Green pentagon represents *attP2*, blue pentagons represent *attP40*, red/blue composite pentagons represent *attP40* carrying indicated insertion. Brown rectangle represents insertion. E) Percentage of nuclei found within clusters for each genotype. Means +/-SEMs are shown. The following calculation was used: ((number of nuclei within clusters / total number of nuclei) x 100). F) Percentage of hemisegments with normal nuclear distribution within muscle 6. Each genotype is represented with a different color in panels D-F with TRiP *spat*/+; *attP2*/+ (green), TRiP *spat*/*attP40* (blue) and TRiP *spat* (red). For each genotype 3 hemisegments per larvae for 6 larvae total were analyzed. ANOVA with Tukey was used for statistical analyses. Scale bar = 50 µm.

### Effects of *attP40* and derivatives on *MSP300* transcript levels

We hypothesized that nuclei clustering in larvae homozygous for *attP40* and derivatives reflected altered expression of at least some of the eleven *MSP300* isoforms (Figure 1). To test this hypothesis, we prepared RNA from whole third instar larvae and performed quantitative RT-PCR (Q-PCR) using primers from two regions of the *MSP300* transcription unit; first, the far 3’ end of *MSP300*, which includes the KASH domain and accounts for eight of eleven annotated transcripts (Xie and Fischer 2008), see Figure 5A, and second, an internal region that is predicted to amplify the *RB* and *RF* transcripts, which lack the KASH domain. Thus, these primers enable analysis of 10 of 11 annotated transcripts (Figure 5A).

**Figure 5.**
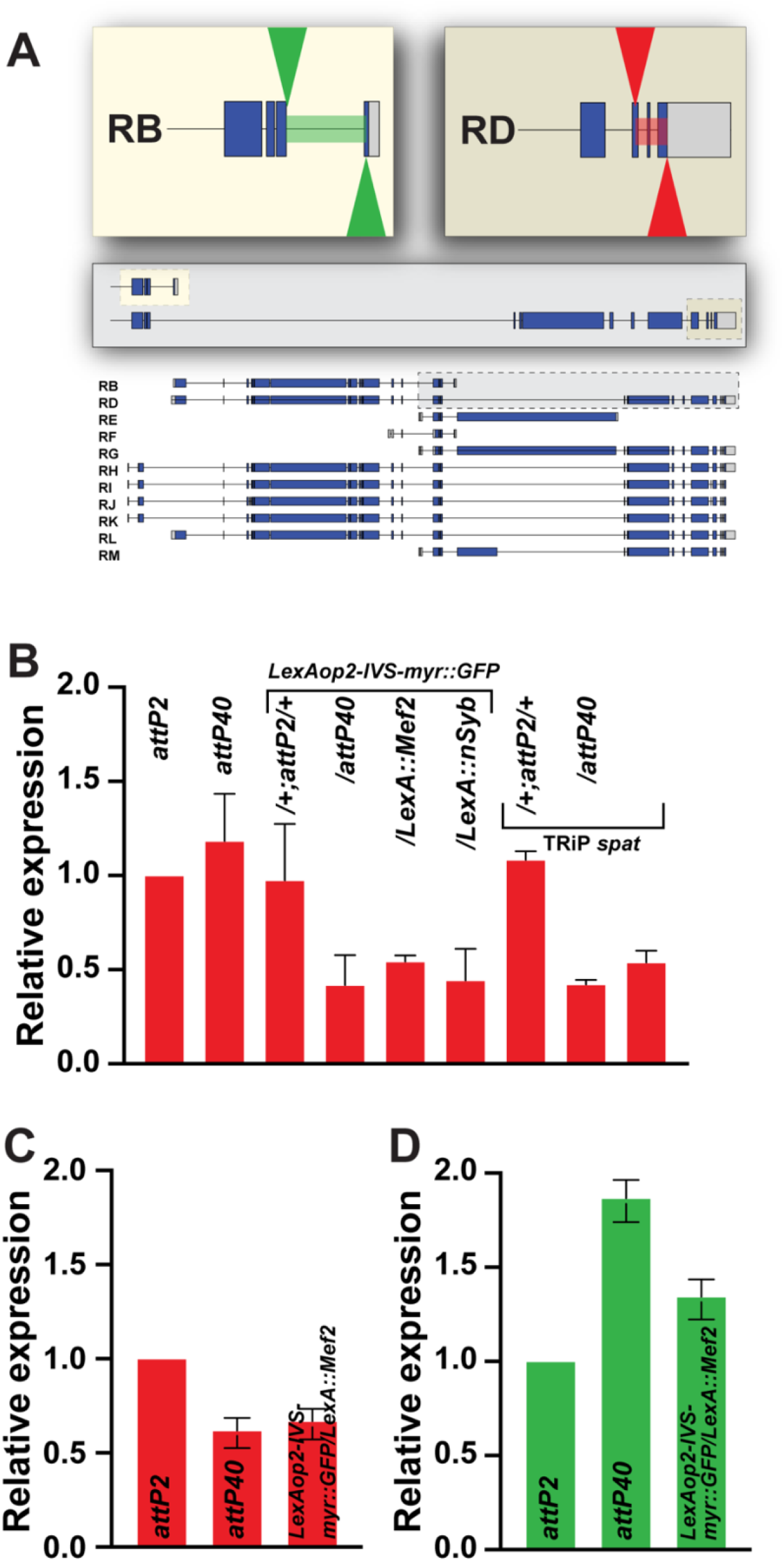
Effect of *attP40* and derivatives on *MSP300* transcript levels. A) Schematic overview of the primer set used in this analysis (green arrowheads for mid region primers, red arrowheads for 3’ primers). Grey inset enhances the mid to 3’end region of transcripts RB and RD. The (light color) inset shows the primer locations used to analyze transcripts RB and RF. The (dark color) inset show the primer locations used to analyze transcripts RD, RG, RH, RI, RJ, RK, RL and RM (KASH-containing transcripts). Annotated transcripts as shown in Figure 1. B) Quantitative RT-PCR (Q-PCR) was used to measure *MSP300* KASH-containing transcript levels, normalized to *attP2*, from whole third instar larvae of the indicated genotypes. C) Q-PCR was used to measure *MSP300* KASH-containing transcript levels, normalized to *attP2*, from 3^rd^ instar larval fillets of the indicated genotypes. D) Q-PCR was used to measure *MSP300* transcripts B and F, normalized to *attP2*, from whole third instar larvae of the indicated genotypes. Red bars represent transcript levels measured with *MSP300* KASH-containing primers and green bars represent transcript levels measured with transcripts B and F primers. Mean +/-SEM was determined from at least three separate biological samples collected from each genotype, and triplicate measures of each sample. The 2^-ΔΔCt^ was employed for this measurement. Each sample contained a mix of 10 whole larvae for (B) and (D) or a mix of 5 larval fillets (C).

We found using the 3’ end primers that *MSP300* transcript levels were decreased about two-fold in larvae homozygous for *attP40* and that contained an insertion in at least one of the *attP40* sites (Figure 5B). These results support the possibility that *attP40* derivatives cause nuclei clustering by decreasing *MSP300* transcript levels. These effects on transcript levels, like effects on nuclear clustering, were recessive, as *MSP300* transcript levels in *LexAop2-IVS-myr::GFP/+* or TRiP *spat/+* larvae were not distinguishable from those from the *attP2* control.

We were surprised to find that larvae homozygous for empty *attP40* exhibited wildtype *MSP300* transcript levels, despite exhibiting a strong nuclear clustering phenotype. However we note that *MSP300* is transcribed in larval fat bodies as well as muscle (Zheng *et al*. 2020); thus our whole larva RNA preparations might not capture *attP40* transcriptional effects specifically in muscle. To address this possibility, we performed Q-PCR on RNA prepared from larval fillets, in which the fat body as well as all other internal organs were removed, leaving only the body wall muscles, underlying epidermis, and cuticle. We found that with this semi-purified muscle tissue as RNA source, empty *attP40* fillets as well as *LexAop2-IVS-myr::GFP/LexA::Mef2* fillets exhibited a ∼two-fold decrease in *MSP300* transcript levels (Figure 5C), similar to what we observed in whole larvae carrying *attP40* insertions (Figure 5B). Thus, muscle *MSP300* transcript levels match muscle Msp300 phenotypes in *attP40* and derivatives. These results also raise the possibility that empty *attP40* might increase levels of certain *MSP300* isoforms in non-muscle tissues.

The primers used in Figure 5B enable amplification of KASH domain isoforms. To analyze transcript levels of the two annotated non-KASH domain isoforms, transcripts *RB* and *RF*, we used the internal primers described above to amplify RNA from whole larvae. We found that unlike the case with the KASH domain transcripts, non-KASH domain transcripts were increased in both empty *attP40* and *LexAop2-IVS-myr::GFP/LexA::Mef2* larvae (Figure 5D). Phenotypic consequences of these altered transcript levels are not clear. Thus, *attP40* can have distinct effects on transcript levels of different isoforms.

### Effects of TRiP insertions into *attP40* on viability and fertility

Many transgenes introduced into *attP40* are RNAi short hairpin sequences from the “TRiP” project cloned into the *Valium20* vector. We noticed some unexpected viability phenotypes, even in the absence of expression, when working with some of these insertions, so we wanted to characterize TRiP insertion viability systematically. However, monitoring TRiP insertions for recessive viability and fertility was problematic for two reasons. First, many TRiP insertions into *attP40* are balanced with *CyO;* this is problematic because the *Cy*^*1*^ “Curly” dominant marker on the *CyO* balancer is unreliable. The Curly wing phenotype is easily suppressed by genetic modifiers as well as the environmental conditions of low temperature and larval crowding (Nozawa 1956); indeed, the FlyBase allele report for *Cy*^*1*^ states that *Cy*^*1*^ “frequently overlaps [wildtype] at 19°C………..some balanced Cy chromosomes pick up suppressors of Cy in stock” (http://flybase.org/reports/FBal0002196.html). Second, the *y*^*+*^ and *v*^*+*^ markers used for the TRiP insertions are each fully dominant, unlike the semi-dominant mini-white marker used on other transgenes. Because of these two features, it is difficult to accurately distinguish flies carrying *CyO* from flies without *CyO* by simple visual inspection.

To address these difficulties, we placed several TRiP insertions in combination with a modified *CyO* balancer upon which the dominant *Tb*^*1*^ “Tubby” transgene had been introduced by P-element mediated transformation (Lattao *et al*. 2011). This *CyO-TbA* balancer also carries the dominant *Star* marker, which confers “Rough eyes” (Kolodkin *et al*. 1994). Both *Tb*^*1*^ and *Star* are completely penetrant. Thus, the use of *CyO-TbA* to balance TRiP insertions enabled us to unambiguously distinguish flies heterozygous from homozygous for each TRiP insertion at the larval, pupal, and adult stages.

From six TRiP insertions tested when homozygous, we found a wide variety of viability and fertility deficits (Table 1 and Table 2). The mostly strongly affected insertions, TRiP *atl* and TRiP *spat*, permitted no homozygous viable adults, although a few pupae homozygous for TRiP *spat* were observed. Flies bearing the TRiP *rop* and *TRiP Mcu* insertions were less strongly affected. Pupae homozygous for either insertion were observed, albeit at a lower frequency than expected, and most (∼80%) failed to eclose. Escaper adults exhibited greatly decreased fertility (Table 2). In particular, none of the males homozygous for TRiP *rop*, and only 2 of 10 males from TRiP *Mcu*, were fertile. Likewise, most females homozygous for TRiP *rop* and TRiP *Mcu* were infertile, and the rare fertile females produced very few progeny. Flies bearing the TRiP *JNK* insertion were even less strongly affected. Adults homozygous for TRiP *JNK* were plentiful (Table 1), and the females displayed wildtype fertility (Table 2). However, males homozygous for TRiP *JNK* appeared to be completely sterile (Table 2). All phenotypes of flies homozygous for TRiP *spartin* appeared similar to wildtype. TRiP *spartin* was the only one of the six TRiP lines tested for which we were able to construct and maintain a homozygous stock.

**Table 1:** Homozygous viability in TRiP lines.

| Genotype | Tubby (T) | Non-Tubby (NT) | NT/Total (%) | Non-Tubby eclosion | Eclosion (%) |
| --- | --- | --- | --- | --- | --- |
| *attP40* | N.A.* | N.A | N.A | 87/103 | 84.5 |
| TRiP *JNK/CyO-Tb<sup>1</sup>* | 331 | 235 | 41.52% | 203/235 | 86.4 |
| TRiP *spat/CyO-Tb<sup>1</sup>* | 285 | 34 | 10.66% | 0/34 | 0 |
| TRiP *Spartin/CyO-Tb<sup>1</sup>* | 186 | 106 | 36.30% | 81/106 | 76.4 |
| TRiP *Mcu/CyO-Tb<sup>1</sup>* | 183 | 83 | 31.20% | 18/83 | 21.7 |
| TRiP *Rop/CyO-Tb<sup>1</sup>* | 353 | 102 | 22.42% | 27/102 | 26.5 |
| TRiP *atl/CyO-Tb<sup>1</sup>* | 310 | 0 | 0.00% | N.A | N.A |

\*Not Applicable

**Table 2:**
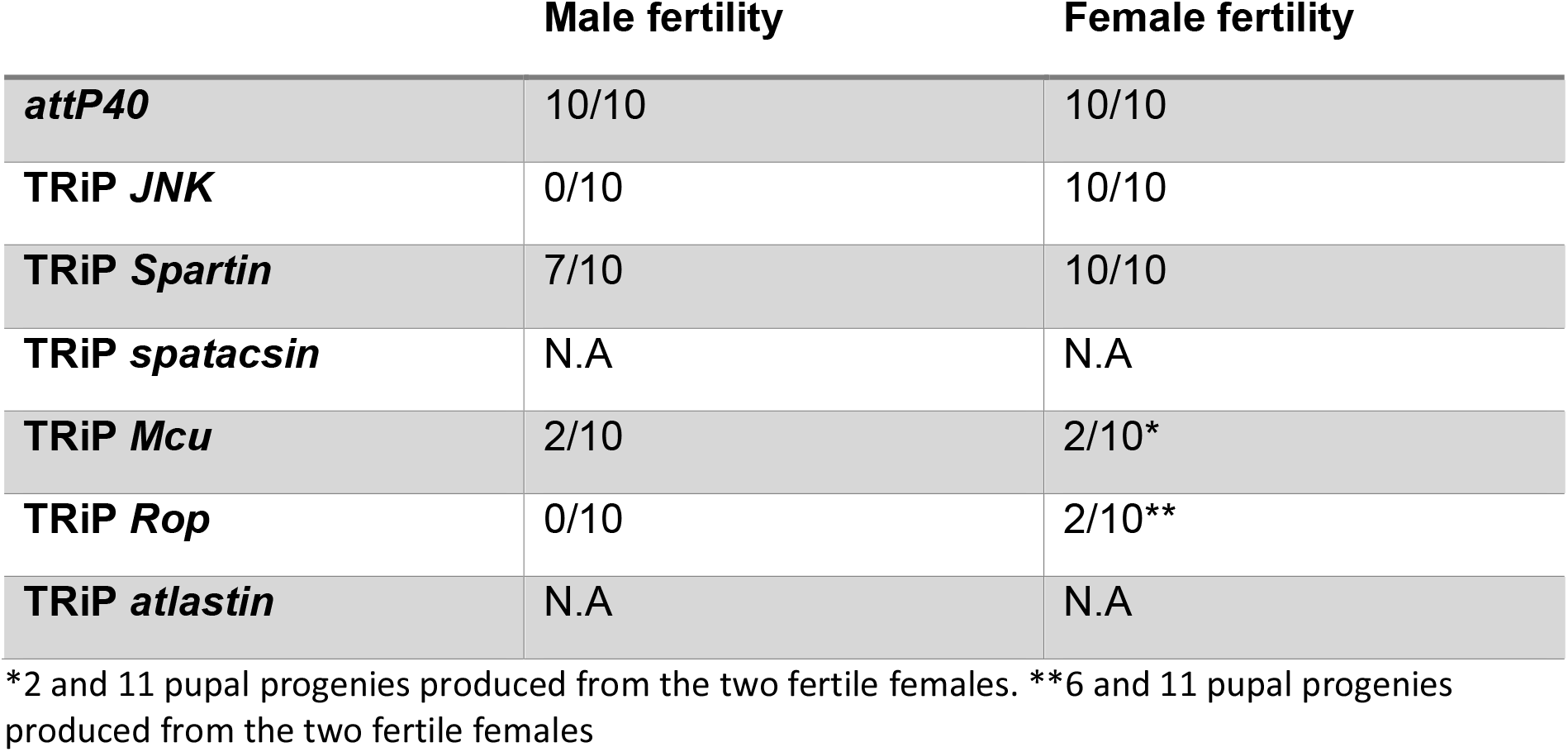
Adult fertility in TRiP lines.

Pupae homozygous for TRiP insertions showed body size defects as well as viability or fertility defects. We did not notice any visible defects in the pupa case and posterior/anterior spiracles in four of our TRiP lines (Figure 6A). Pupae homozygous for *spat* usually failed to evert their anterior spiracles (Figure 6B), a phenotype previously observed in mutants defective in ecdysone production (Mirth *et al*. 2005) or larval muscle function (van der Graaf *et al*. 2021). Although pupal length in these homozygous pupae was not significantly different from the control (pupae homozygous for empty *attP40*), pupal width in these homozygous pupae was significantly decreased (Figure 6C-D). The TRiP *Mcu* insertion had the strongest effect on pupal width.

**Figure 6:**
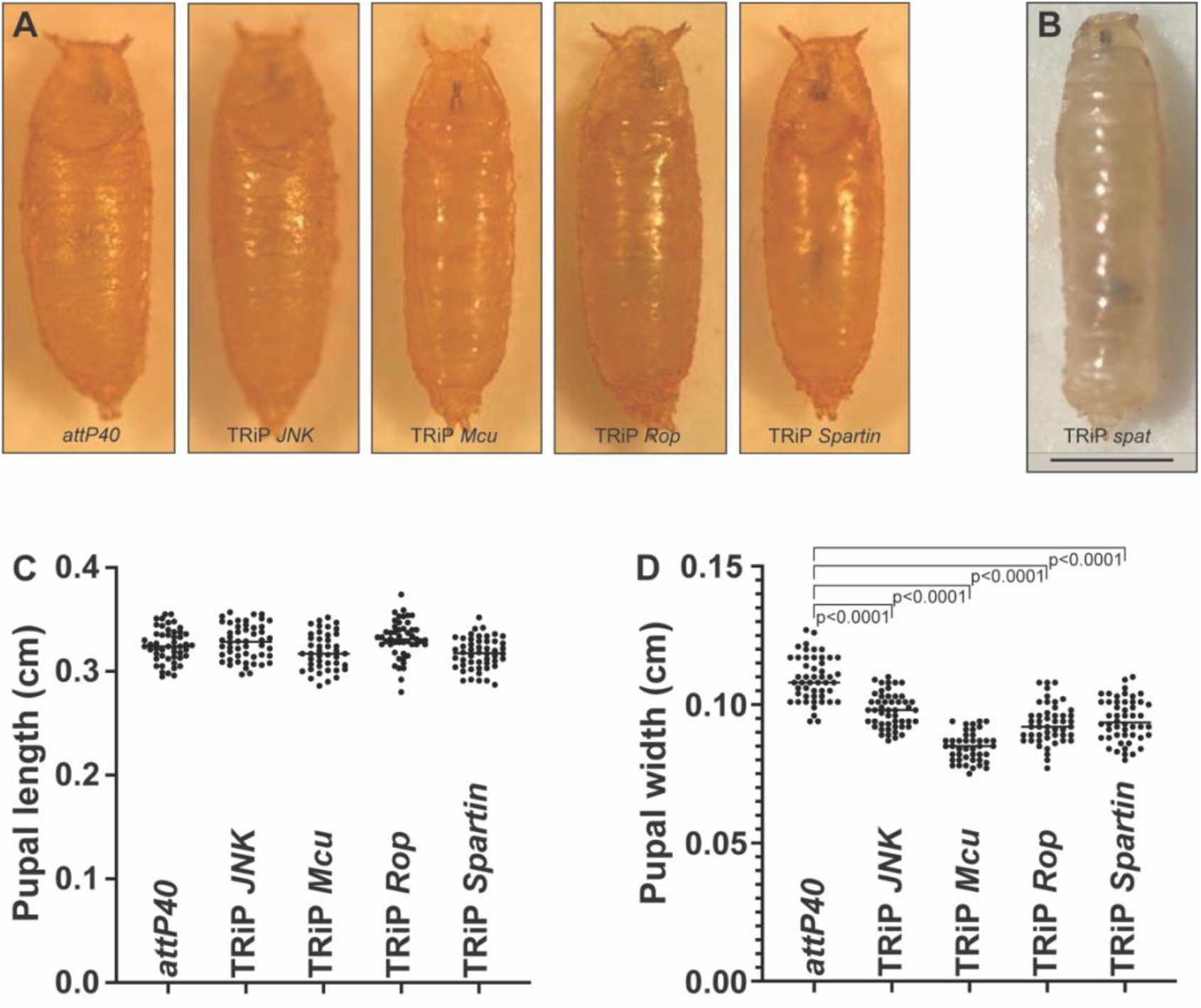
Effects of homozygous TRiP insertions on pupal length and width. A) Visual representation of pupae homozygous for *attP40* and TRiP *JNK, Mcu, Rop* and *Spartin*. B) Visual representation of an early pupa homozygous for TRiP *spat*. C) Pupal length measurements for the genotypes shown in (A). D) Pupal width measurements for the genotypes shown in (A). For pupa length and width analysis, a One-way ANOVA with Bunnett post hoc test was performed. Scale bar=1 mm.

In an attempt to separate the recessive lethality of the TRiP *atl* chromosome from the *attP40* insertion, we performed two rounds of free recombination between TRiP *atl* and another stock with no lethal mutations on chromosome II. However, we were unable to separate the lethality from the TRiP *atl* insertion, indicating that the lethal allele of TRiP *atl* is at or close to the insertion site.

We also tested the viability and fertility of the flies carrying the identical hairpin sequence found in TRiP *atl* but introduced into *attP40* via a different vector. In particular, we introduced the TRiP *atl* hairpin sequence into pJFRC19-13XlexAop2, and then introduced this plasmid into *attP40* with ϕC31-mediated recombination. We found that flies bearing this *Aop-atlRNAi* insertion were fully viable and fertile. Therefore, the phenotype conferred by the hairpin sequence from TRiP *atl* is affected by genetic context.

### Dominance and recessiveness of *attP40* and derivatives

Data shown in Figures 2-5 indicate that *attP40* and derivatives are recessive for nuclear clustering and *MSP300* transcription phenotypes. However, in certain genetic backgrounds, we have found that *attP40* and derivatives can have dominant effects on clustering. In particular, we introduced the pan-neuronal *Gal4* driver *nSyb-Gal4* into flies heterozygous for either empty *attP40*, or the TRiP *atl* insertion into *attP40*. We found that the *nSyb-Gal4* background did not have a significant effect on larval muscle nuclear clustering in empty *attP40/+* (Figure 7); however, TRiP *atl/+* larvae carrying *nSyb-Gal4* displayed a nuclear clustering phenotype significantly different from both empty *attP40/+* and empty *attP40/+; nSyb-Gal4* (Figure 7). Therefore, *attP40* derivatives can confer dominant effects in certain genetic backgrounds. In addition, these results support the conclusion that the TRiP *atl* insertion into *attP40* confers a stronger MSP300 phenotype than empty *attP40*.

**Figure 7.**
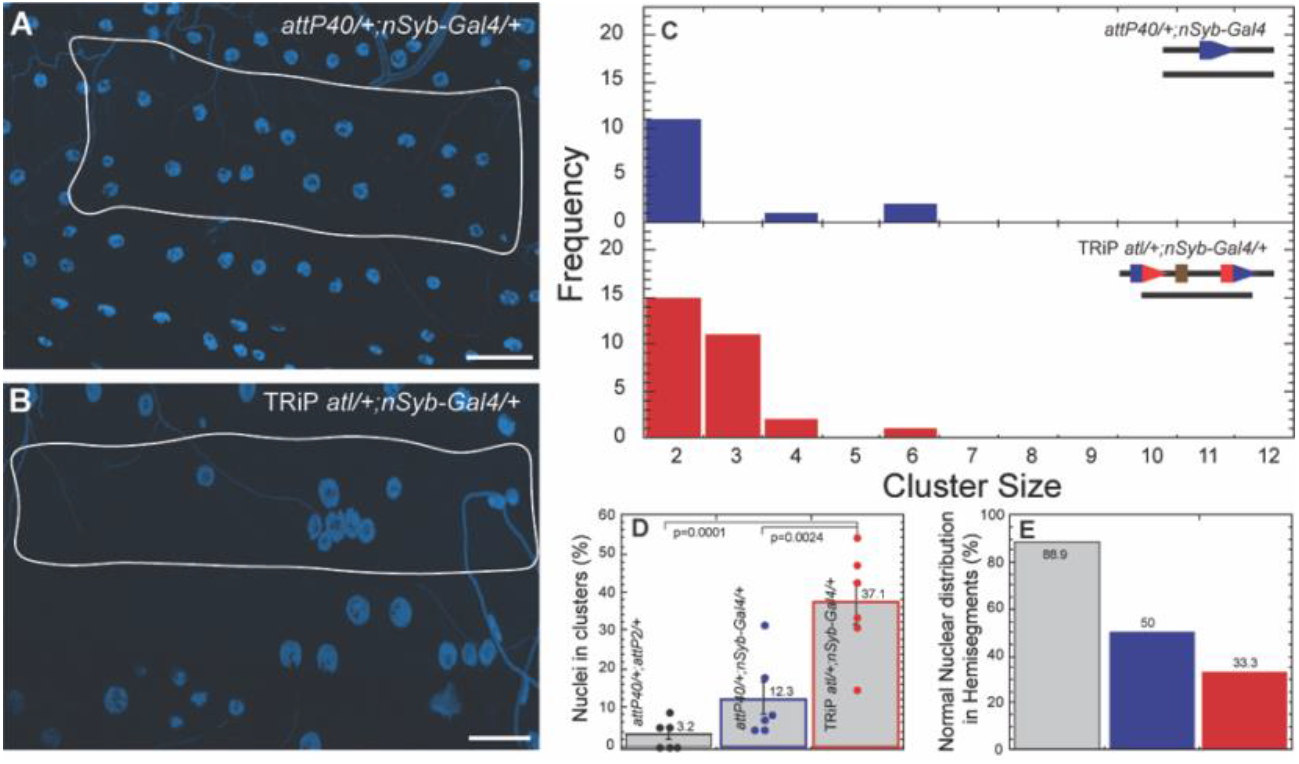
TRiP *atl* can have dominant effects on nuclear clustering in some genetic backgrounds. A-B) Representative images of nuclei position within muscle 6 for *attP40*/+; *nSyb-Gal4*/+ (A) and TRiP *atl*/+; *nSyb-Gal4*/+ (B). Nuclei are labeled with DAPI (blue). Muscles are outlined with solid white lines. C) Frequency distribution of the number of nuclei within each cluster within muscle 6 for each genotype indicated in upper right. Lines in upper right show genotype of each chromosome II homologue. Blue pentagons represent *attP40*, red/blue composite pentagons represent *attP40* carrying indicated insertion. Brown rectangle represents insertion. D) Percentage of nuclei found within clusters for each genotype. Means +/-SEMs are shown. The following calculation was used: ((number of nuclei within clusters / total number of nuclei) x 100). E) Percentage of hemisegments with normal nuclear distribution within muscle 6. Each genotype is represented with a different color in panels C-E with *attP40*/+; *attP2*/+ (grey), *attP40*/+; *nSyb-Gal4*/+ (blue) and TRiP *atl*/+; *nSyb-Gal4*/+ (red). For each genotype 3 hemisegments per larvae for 6 larvae total were analyzed. ANOVA with Tukey was used for statistical analyses. Scale bar = 50 µm.

## Discussion

### Effects of the *attP40* docking site and derivatives on *MSP300*-mediated phenotypes

The *attP40* docking site for Drosophila transgene integration is widely used for ϕC31-mediated transgene integration. *attP40* lies within the transcription unit of *MSP300*, which encodes the Drosophila orthologue of Nesprin-1; this observation raises the possibility that *attP40* is an *MSP300* insertional mutation. Here we show that *attP40*, either alone or containing any of several specific transgene insertions, causes phenotypes similar to *MSP300* mutations, such as larval muscle nuclear clustering and viability deficits. In addition, *attP40-*containing larvae exhibit an approximately two-fold decrease in *MSP300* transcript levels of certain isoforms. The effect on nuclear clustering and viability varies depending on the precise transgene introduced into *attP40*. We conclude that *attP40* is an insertional mutation for *MSP300*.

### Additional *MSP300*-dependent and independent phenotypes conferred by *attP40*

Reports from other investigators are now appearing in which phenotypic consequences of *attP40*-bearing flies are described. In a Ph.D. thesis, Cypranowska (2020) reported that *attP40* decreases *MSP300* transcription and also confers abnormal synapse function at the larval neuromuscular junction. Many of these phenotypes are similar to those conferred by loss of the muscle glutamate receptor *gluRIIA*; this observation is significant because it was previously reported that gluRIIA localization is affected by MSP300 (Morel *et al*. 2014). These observations raise the possibility that *attP40* regulates synaptic transmission via MSP300-dependent effects on gluRIIA.

Other investigators have reported other *attP40*-dependent phenotypes. For example, (Groen *et al*. 2022) reported that *attP40* decreases transcription of *ND-13A*, the gene immediately centromere-distal from *attP40* and which encodes a component of mitochondrial complex I. Further, they suggest that this decreased *ND-13A* transcription might mediate the resistance to the chemotherapy agent cisplatin observed in *attP40* flies (Groen *et al*. 2022). More recently, (Duan *et al*. 2022) reported that *attP40* alters neuronal architecture of the olfactory glomerulus (Duan *et al*. 2022), a phenotype that appears to be MSP300-independent. Taken together, these results indicate that *attP40* can confer a variety of phenotypes in a variety of tissues by affecting transcription of at least two genes.

### *attP40* phenotypes can be dominant or recessive depending on genetic background

We tested if *attP40* and derivatives containing insertions into *attP40* were dominant or recessive for the nuclear clustering and *MSP300* transcription phenotypes. We found that in most cases, the effects of *attP40* and derivatives were recessive. For example, the neutral reporter *LexA-myr-GFP* or the TRiP *spat* insertion within *attP40* each conferred severe nuclear clustering and decreased *MSP300* transcription when homozygous or in combination with empty *attP40*, but not when *attP40* was absent, from the other homologue. These observations indicate that *attP40-*dependent phenotypes are recessive. However, in other genetic backgrounds, we detected dominant effects of *attP40* derivatives. In particular, the TRiP *atl* insertion within *attP40* conferred a dominant nuclear clustering phenotype in a genetic background containing the neuronal Gal4 driver *nSyb-Gal4*. Similarly, (Cypranowska 2020) reported that *attP40* conferred a dominant 2.7-fold decrease in *MSP300* transcript levels in the presence of the motor neuron Gal4 driver *OK6*. We conclude that the phenotypes of *attP40* and derivatives can be dominant or recessive depending on genetic background.

### Effects of specific transgene insertions into *attP40* on mutant phenotypes

We tested if the nucleotide sequence of specific transgenes inserted into *attP40* would affect mutant phenotypes. First, we found that transgenes from either the *LexA* or the *Gal4* regulatory systems were capable enhancing the nuclear positioning phenotype and *MSP300* transcript phenotypes to moderate degrees. Second, we tested if transcription of inserted transgenes in muscle would affect mutant phenotypes. In particular, we compared nuclear positioning in larvae expressing *LexA-myr-GFP* in neurons vs. muscle and found that muscle expression modestly increased severity of the nuclear clustering phenotype but did not affect *MSP300* transcript levels. These results indicate that muscle transcription of inserted transgenes is not necessary for mutant phenotype, but we are unable to rule out the possibility muscle transcription could contribute to severity of mutant phenotype.

### TRiP insertions into *attP40*: extremely variable degrees of recessive lethality and sterility

Of the 16,503 lines carrying *attP40* maintained at the Drosophila stock center (Bloomington, IN), 8,884 contain TRiP (short hairpin sequences for RNAi) insertions, mostly in the *Valium 20* vector. In studies of six of these lines, we observed a range of recessive phenotypes, from full lethality to full (or partial) sterility to ∼wildtype. This extreme variability in phenotypes among the TRiP lines was unexpected, as each lines contains inserts with identical sequences, with the exception of the sequence of the short hairpin itself. These results suggest either that phenotypic strength is affected unexpectedly strongly by the precise nucleotide sequence of the hairpin, or alternatively, that these TRiP lines have accumulated genetic modifiers at an unusually high rate, and that these modifiers have variable effects on phenotype. These two possibilities are not mutually exclusive.

Because of the wide variety of phenotypes exhibited by flies carrying distinct TRiP insertions, we wondered if we could predict phenotypes conferred by specific TRiP lines from information presented at the Bloomington Drosophila Stock Center (BDSC, https://bdsc.indiana.edu). We noticed that in the description of the six TRiP lines tested, five (TRiP *atl*, TRiP *spat*, TRiP *rop*, TRiP *Mcu*, and TRiP *JNK*) contained the phrase “May be segregating CyO”, or a related phrase. These five lines are the same ones for which we were unable to maintain a homozygous line. The sixth line (TRiP *spartin*), for which we were able to maintain a homozygous line, lacked this phrase. This correspondence raises the possibility that researchers might be able to distinguish homozygous viable and fertile TRiP lines from others by the absence/presence of this phrase. As of the most recent report (from 2019), the BDSC reports that 31.46% of the 8884 TRiP lines in *attP40* (2795 total) are listed as homozygous, with the rest as “CyO”, “CyO fl” or “CyO mix”.

## Conclusions

We have shown *attP40* and derivatives containing insertions confer *MSP300* mutant phenotypes and can decrease *MSP300* transcript levels. Regardless of the mechanism underlying the variety of phenotypes conferred by various TRiP insertions within *attP40*, investigators should be aware that these insertions might confer phenotypes that are difficult to predict and might manifest in a variety of ways. Going forward, investigators should use caution when interpreting data resulting from flies containing *attP40*, especially in muscle tissues.

## Acknowledgments

The authors would like to thank the Bloomington Stock Center (BDSC) for providing all the fly lines used in this study. We are grateful to Annette Parks (BDSC) for communicating information on BDSC fly stock holdings and to Alekhya Gurram for assistance with Q-PCR experiments. This work was done in part using resources of the Rice University Shared Equipment Authority (https://research.rice.edu/sea/). We thank Budi Utama for help with light microscopy and image quantitation. This study is funded by grants R01 NS102676 and R21 NS111340 to MS and JAM.

The authors declare that they have no conflicts of interest with the contents of this article.

## Author Contributions

KvdG – Collected the data, designed the analysis, performed the analysis, wrote manuscript, edited the manuscript; SS – performed the analysis; PS – performed the analysis; MS – Conceived and designed the analysis, edited the manuscript; JAM – Conceived and designed the analysis, edited the manuscript.

## Notes

### Competing Interest Statement

The authors have declared no competing interest.

### Summary of Updates

We have reworked all figures to show the clustering phenotype more clearly. Added additional Q-PCR analysis of MSP300. New data added of the recessive lethality and/or sterility of several TRiP insertions.

